# Chronic Bilateral Cochlear Implant Stimulation in Early-Deaf Rabbits Restores Neural Binaural Sensitivity

**DOI:** 10.1101/803627

**Authors:** Woongsang Sunwoo, Bertrand Delgutte, Yoojin Chung

**Author notes:** Correspondence to: Yoojin Chung, Eaton-Peabody Laboratories, Massachusetts Eye & Ear Infirmary, Boston, MA 02114, USA. Conflict of interest: None.

## Abstract

Cochlear implant (CI) users with a pre-lingual onset of hearing loss show poor sensitivity to interaural time differences (ITD), an important cue for sound localization and speech reception in noise. Similarly, neural ITD sensitivity in the inferior colliculus (IC) of neonatally-deafened animals is degraded compared to animals deafened as adults. Here, we show that chronic bilateral CI stimulation during development can partly reverse the effect of early-onset deafness on ITD sensitivity. The prevalence of ITD sensitive neurons was restored to the level of adult-deaf rabbits in the early-deaf rabbits that received chronic stimulation with wearable bilateral sound processors during development. In contrast, chronic CI stimulation did not improve temporal coding in early-deaf rabbits. The present study is the first report showing functional restoration of ITD sensitivity with CI stimulation in single neurons and highlights the importance of auditory experience during development.

## Introduction

Although cochlear implantation (CI) provides effective treatment for profound deafness, both adult and pediatric CI users still face challenges understanding conversations in everyday noisy acoustic environments. Normal hearing listeners benefit from using binaural cues for sound localization and understanding speech in noise. A growing number of CI users, including children, receive implants in both ears to benefit from these binaural cues as well. However, binaural performance of bilateral CI users is poorer than that of normal hearing listeners, particularly for tasks that requires sensitivity to interaural time differences (ITD) (Salloum et al., 2010; van Hoesel, Jones, & Litovsky, 2009; van Hoesel & Tyler, 2003). As a result, bilateral CI listeners’ abilities to understand speech in everyday acoustic environments and localize sound sources are worse than normal (Litovsky, Parkinson, & Arcaroli, 2009; Murphy & O’Donoghue, 2007). CI users’ sensitivity to ITD is generally poorer than that of normal hearing listeners and highly dependent on stimulus parameters such as pulse rate and intensity (Kan & Litovsky, 2015; Laback, Egger, & Majdak, 2015). This problem is especially acute in patients who experience early-onset hearing loss (Litovsky, Jones, Agrawal, & van Hoesel, 2010). CI users with pre-lingual onset of hearing loss who received CIs as adults cannot perceive ITDs or perform very poorly in ITD discrimination tasks. Single neurons in the central auditory pathway show similar limitations in how they encode bilateral CI stimulation. Degraded neural ITD sensitivity with bilateral CI stimulation has been observed in the auditory midbrain (Hancock, Noel, Ryugo, & Delgutte, 2010) and auditory cortex (Tillein et al., 2010) of congenitally deaf cats under anesthesia, and auditory midbrain of neonatally-deafened rabbits in an unanesthetized preparation (Chung, Buechel, Sunwoo, Wagner, & Delgutte, 2019).

Whether chronic auditory stimulation through bilateral CIs can restore ITD sensitivity in central auditory neurons is unclear. Children who received bilateral CIs at an early age show some improvement in perception of binaural cues, although their ITD sensitivity remains poorer than that of normal hearing children (Gordon, Deighton, Abbasalipour, & Papsin, 2014). Similarly, evoked potential studies in congenitally deaf children who received simultaneous bilateral CIs show that auditory experience with current clinical devices does not fully reverse effects of early deafness (Easwar, Yamazaki, Deighton, Papsin, & Gordon, 2017). Anatomical studies in congenitally deaf cats have shown that chronic bilateral CI stimulation during development can restore a more normal distribution of inhibitory inputs to neurons in the medial superior olive (MSO), one of the brainstem nuclei where ITD processing first takes place (Tirko & Ryugo, 2012). To what extent neural ITD sensitivity can be restored with chronic bilateral CI stimulation is unknown.

We developed a novel preparation of neonatal deafening, early-implantation, and chronic stimulation with wearable bilateral processors in rabbits. With this preparation, we investigated whether chronic bilateral CI stimulation during development can reverse the effect of early-onset deafness on neural ITD sensitivity and temporal coding in the auditory midbrain. Neurophysiological recordings were made from unanesthetized animals because the upper limit of neural temporal coding and the pulse rate dependence of ITD sensitivity of auditory midbrain neurons better match perceptual results from human CI subjects in unanesthetized animals than in anesthetized animals (Chung, Hancock, & Delgutte, 2016; Chung, Hancock, Nam, & Delgutte, 2014). We found that chronic bilateral CI stimulation during development can partly reverse the degradation in neural ITD sensitivity resulting from early-onset deafness. The effect is most pronounced in response to high-rate stimulation. However, there was no reversal in the effect of early-onset deafness on temporal coding.

## Results

We recorded from 132 single units in the inferior colliculus (IC) of four rabbits that were neonatally deafened and provided chronic stimulation with bilateral CIs starting at the age of 2 months (early-deaf, chronically stimulated, ED-CS). We used an experimental “Fundamental Asynchronous Stimulus Timing” (FAST) strategy specifically designed to deliver ITD cues (Smith, 2010, 2014). Neural ITD sensitivity and temporal coding metrics were compared to previous reported data from 117 neurons in three neonatally deafened and unstimulated rabbits (early-deaf, unstimulated, ED-US) (Chung et al. 2019) and 180 neurons in four rabbits that were deafened as adults (adult-deaf, AD) (Chung et al. 2016) (Figure 1). Detailed history of age at implantation and recording sessions for all groups of rabbits is given in Table 1.

**Figure 1.**
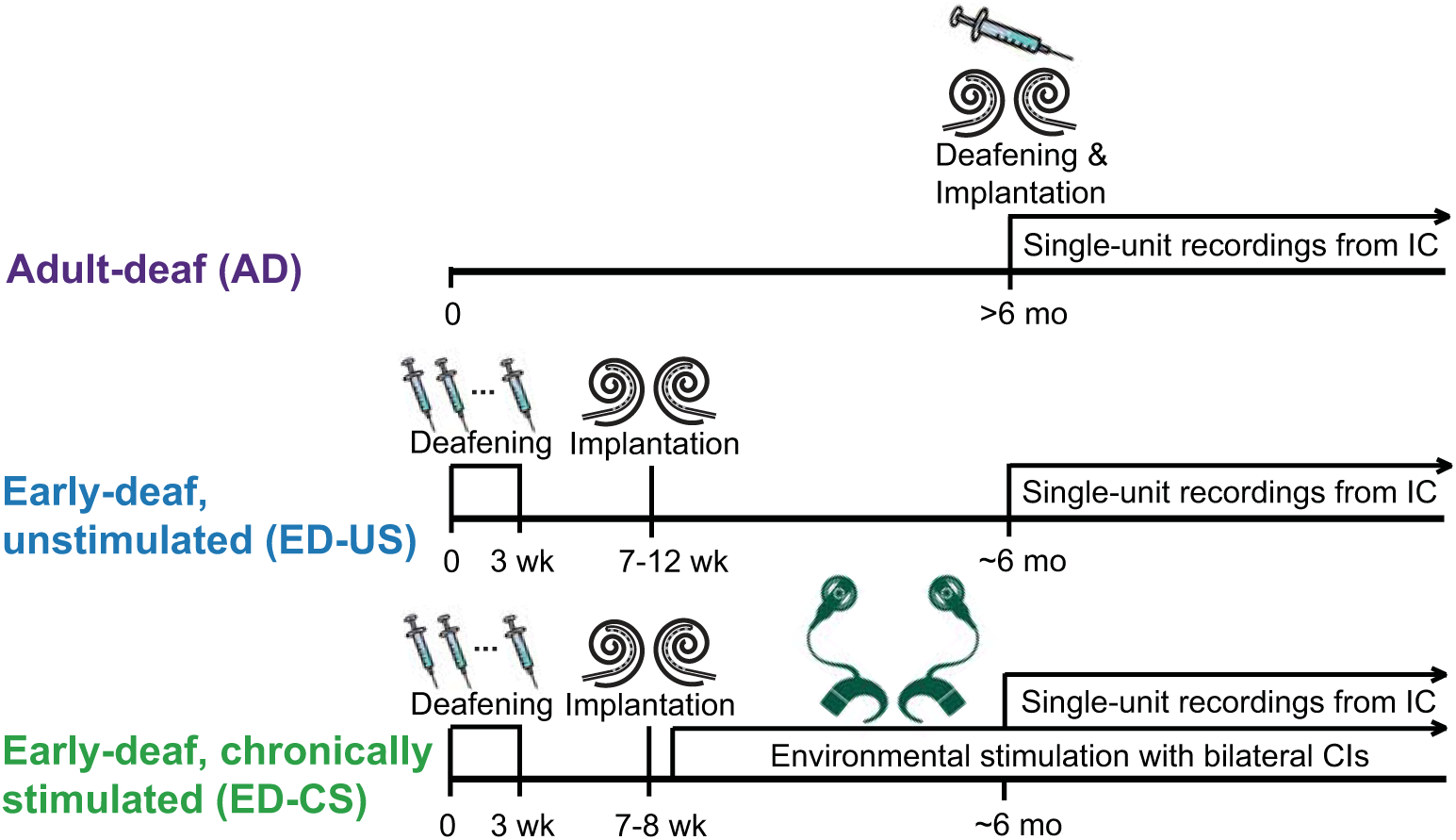
Timeline and order of procedures in three animal groups. AD rabbits were deafened and implanted as adults. ED-US rabbits and ED-CS rabbits were deafened as neonates and implanted at ∼2 months of age. ED-CS rabbits received chronic stimulation immediately after implantation.

**Table 1.**
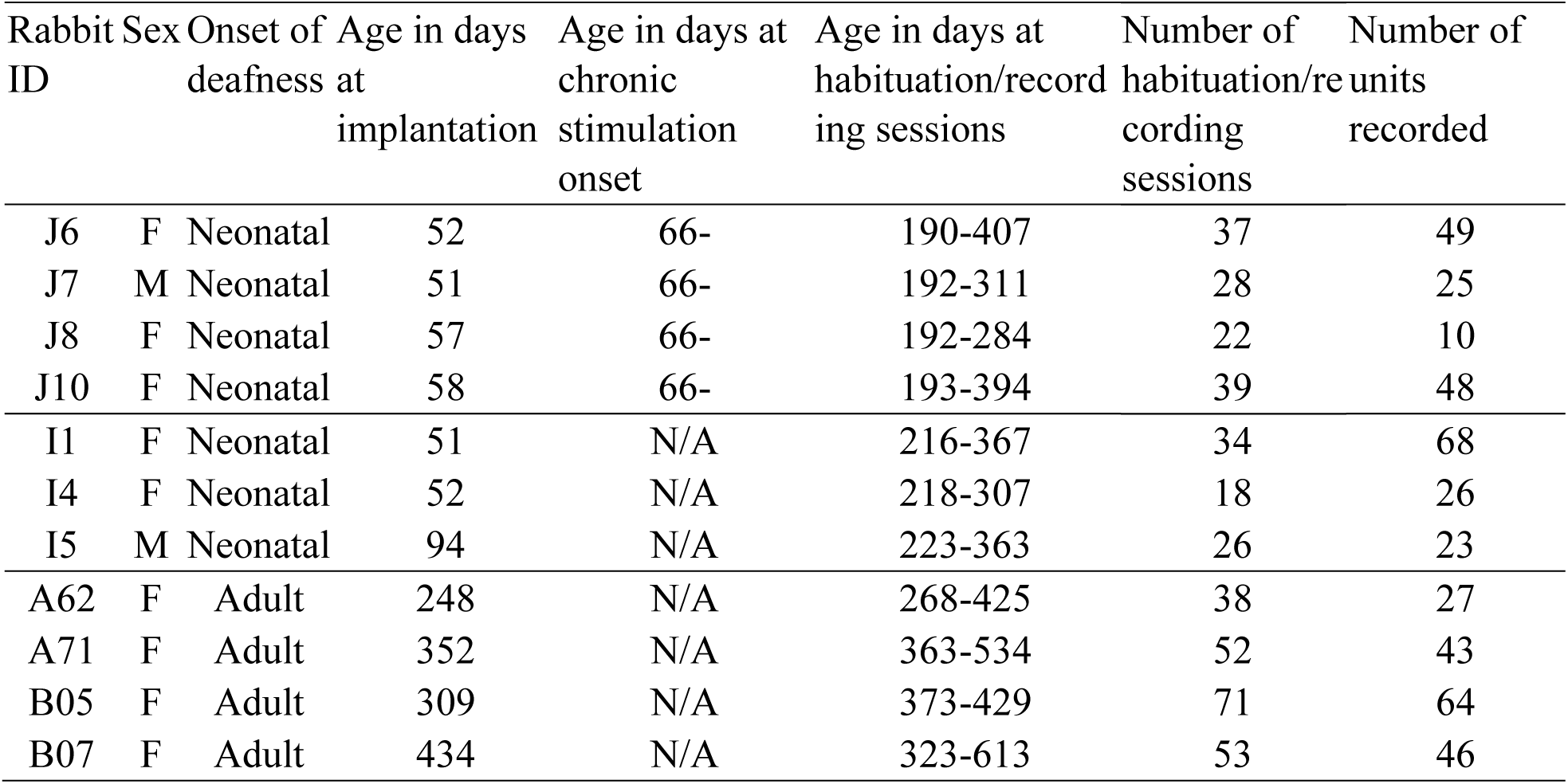
Summary of deafness, auditory experience, and recording history for each animal. For adult deafened rabbits, the onset of deafness is the same as the age at implantation (i.e., all adult-deafened rabbits were deafened during the CI surgery).

### Restoration of ITD sensitivity in early-deaf animals with bilateral CI stimulation

Neural ITD sensitivity of single IC neurons were characterized by measuring responses to periodic pulse trains with ITDs ranging from −2000 to +2000 μs and pulse rates from 20 to 640 pps. Figure 2A-C show temporal response patterns to 160-pps pulse trains with varying ITDs obtained in an example neuron from an ED-US rabbit, an ED-CS, and an AD rabbit. The neuron from the ED-US rabbit shows responses primarily at the onset of the stimulus. We used the ITD signal-to-variance ratio (STVR), an ANOVA-based metric that ranges from 0 (no sensitivity) to 1 (perfectly reliable tuning) to quantify neural ITD sensitivity (Chung et al., 2016; Hancock et al., 2010). The ITD tuning curve from the ED-US rabbit does not show significant ITD sensitivity (Figure 2D). In contrast, the neuron from the rabbit that received bilateral CI stimulation during development (ED-CS) shows high rate of firings at the stimulus onset to contralateral-leading ITDs (>0) and sustained response to 0-400 µs. This peak-shaped ITD sensitivity is clear in the rate-ITD curve in Figure 2D with significant ITD sensitivity based on ITD STVR. The neuron from the AD rabbit also shows contralateral-preferring ITD sensitivity with sustained firing near 400 µs (Figure 2C) and significant ITD sensitivity (Figure 2D). The neuron from the ED-CS rabbit and the neuron from the AD rabbit show comparable ITD sensitivity.

**Figure 2.**
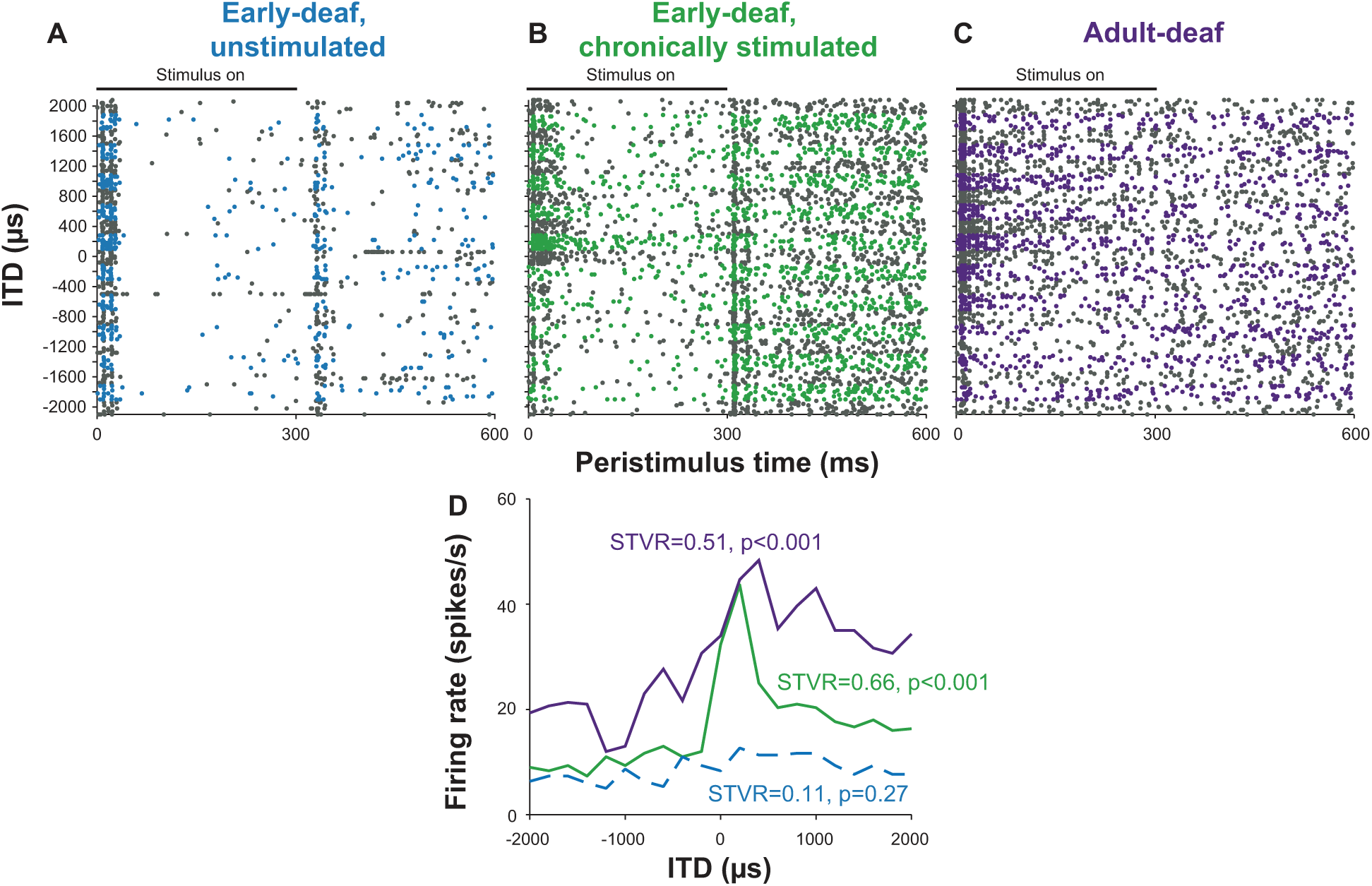
Temporal discharge pattern (dot rasters) as a function of ITD for example neurons from an early-deaf and unstimulated rabbit (A), an early-deaf and stimulated rabbit (B), and an adult-deaf rabbit (C). Alternating colors indicate blocks of stimulus trials at different ITDs. D, Firing rate vs. ITD curves for the three example neurons. Solid lines indicate statistically significant ITD sensitivity based on ANOVA (*p*<0.01).

The ITD sensitivity of IC neurons to bilateral CI stimulation degrades with increasing pulse rates, consistent with limitations in perceptual ITD sensitivity in human CI listeners (Chung et al., 2016). This rate-dependent degradation of ITD sensitivity is more severe in early-deafened and unstimulated animals than in adult-deafened rabbits (Chung et al., 2019). We compared the dependence of ITD sensitivity on pulse rates to investigate whether bilateral CI stimulation during develop can restore ITD sensitivity over a wide range of pulse rates. Figure 3A-C show ITD tuning curves in example units from an ED-US animal, an ED-CS animal, and an AD animal (same neurons as in Figure 2). The neuron from an ED-US rabbit (Figure 3A) shows a weak preference to contralateral-leading ITDs at low pulse rates (40-80 pps) but little or no ITD sensitivity at high pulse rates. In contrast, the neuron from an ED-CS rabbit (Figure 3B) shows sharper ITD tuning with a peak near 400 µs for all pulse rates tested. The neuron from an AD rabbit also shows broad ITD tuning with a preference for contralateral-leading ITDs over a wide range of frequencies (Figure 3C). The ITD STVR (Figure 3D) was higher and statistically significant for most rates tested in the ED-CS neuron (all pulse rates tested) and AD neuron (except for 320pps) than in the ED-US neuron (40-80 pps).

**Figure 3.**
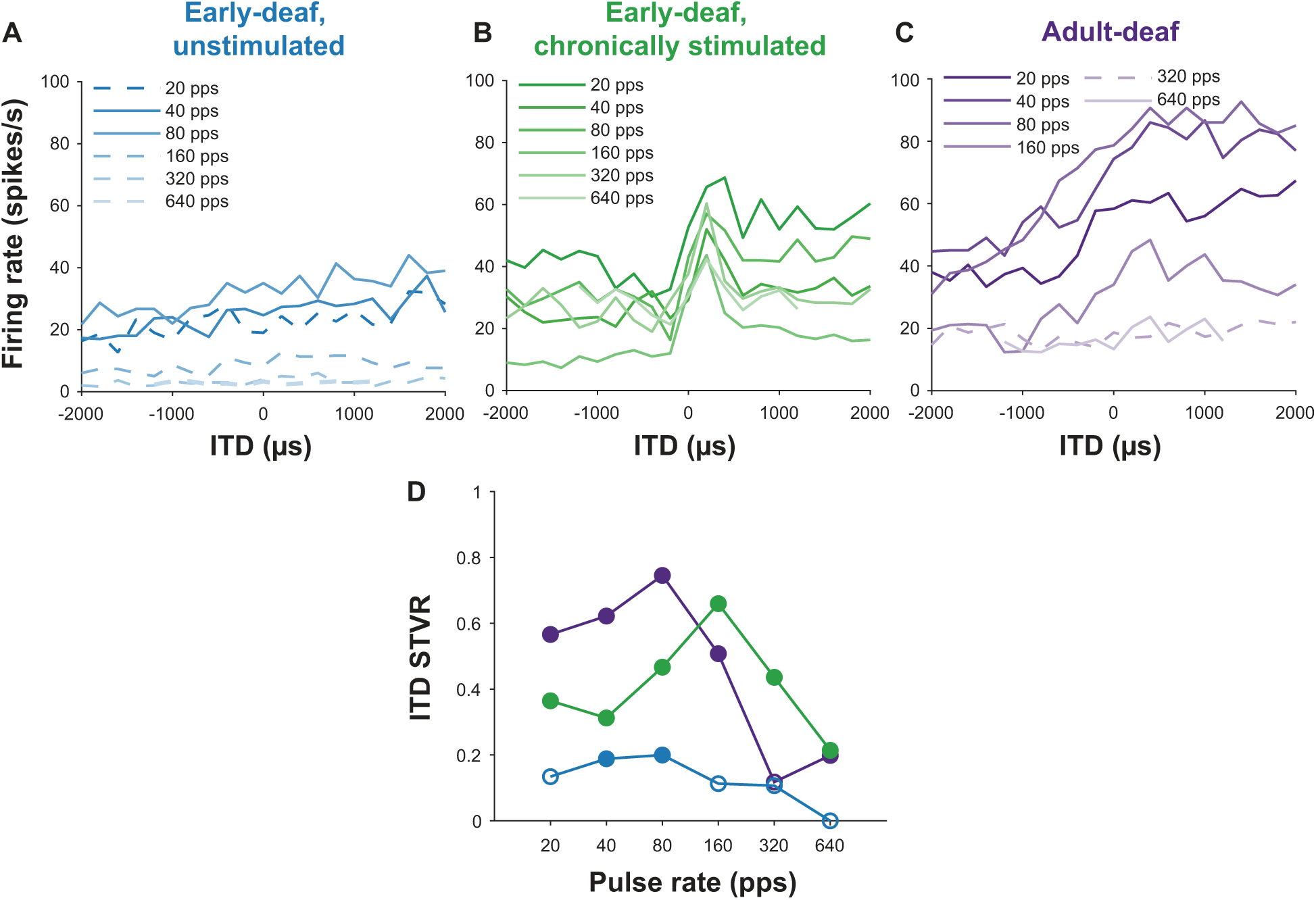
A-C, Firing rate vs. ITD curves for a range of pulse rates for the same three example neurons as in Fig. 1. D, ITD STVR as a function of pulse rate for the three example neurons. Solid lines (A-C) and filled dots (D) indicate statistically significant ITD STVR (*p*<0.01).

The restoration of ITD sensitivity over a wide range of pulse rates seen in example neurons in Figure 3 is representative of the corresponding neural populations. For all pulse rates tested, more neurons from the ED-CS group were sensitive to ITD, as measured by the STVR, compared to the ED-US group (Figure 4A). Overall the fraction of ITD sensitive neurons in the ED-CS group is comparable to the AD group at most rates. This analysis was limited to units that were tested for at least two pulse rates. A two-way ANOVA on the arcsine-transformed fraction of ITD sensitive units showed significant effects of both pulse rate (*F*(5, 17)=25.49, *p*<0.001) and deafness group (*F*(2, 17)=25.08, *p*<0.001). Post hoc paired comparisons (Tukey-Kramer) showed that the fraction of ITD sensitive neurons was significantly lower in the ED-US group compared to the each of the other two groups (ED-CS: *p*=0.0006, AD: p=0.0002), whereas there was no difference between the ED-CS and AD groups (*p*=0.5751). The mean ITD STVR was higher for ED-CS group for most rates (>20 pps) compared to ED-US group but lower for all pulse rates tested compared to the AD group (Figure 4B). The largest effect of chronic stimulation during development is seen for pulse rates above 80 pps. A two-way ANOVA on the arcsine-transformed STVRs showed significant effects of both pulse rate (*F*(5, 1304)=20.72, *p*<0.001) and deafness group (*F*(2, 1304)=14.27, *p*<0.001) but no interaction (*F*(10, 1304) = 0.6, *p*=0.81). Post hoc paired comparisons (Tukey-Kramer) showed that the ITD STVR was significantly higher in ED-CS group compared to ED-US (*p*=0.0032), but significantly lower compared to the AD group (*p* =0.0098).

**Figure 4.**
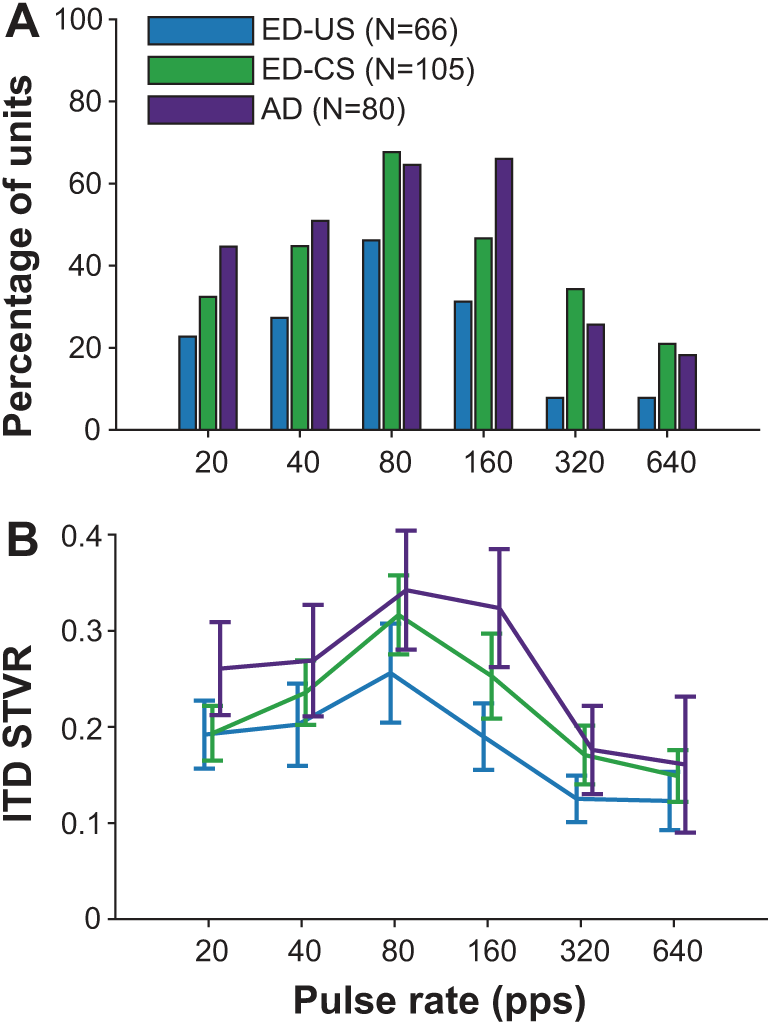
A, Fraction of ITD-sensitive neurons as a function of pulse rate for early-deaf and unstimulated, early-deaf and stimulated, and adult-deaf animals. B, Mean ITD STVR vs. pulse rate for the three animal groups. Error bars indicate ± 2 S.E., which corresponds to a 95 % confidence interval for normal distributions.

We compared the shapes of ITD turning curves from the three groups of rabbits. The ITD tuning curves were fit to a flexible template to classify their shapes into four categories: monotonic, peak, trough, and other (unclassified). Only units that showed significant ITD sensitivity based on the STVR for at least one pulse rate below 160 pps were used for the shape type classification. For units that showed significant ITD sensitivity for multiple pulse rates, the ITD tuning curve with the highest STVR was used for shape classification. The relative incidences of four shapes of ITD tuning curves are compared in Figure 5A. The relative incidences of four types are similar in all three animal groups although there are slightly more unclassified types in the two early-deaf groups (χ^2^(6, N=229)=2.4605, *p*=0.8729).

**Figure 5.**
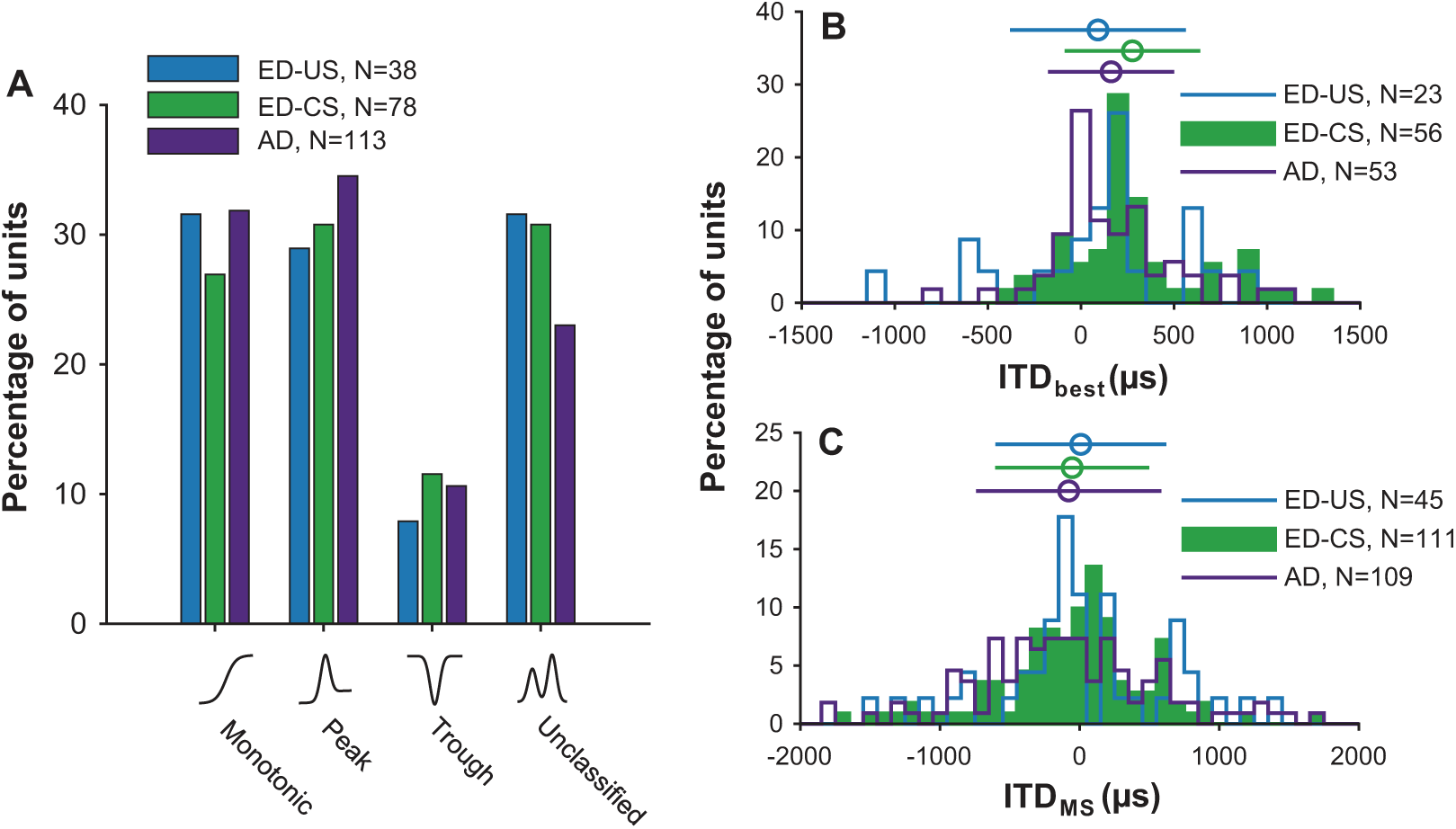
A, Distribution of ITD shape types for the three animals groups. Distribution of best ITDs (B) and ITD_MS_ (C) for the three animal groups. Error bars above each histogram represent means ± 1 S.D.

Two ITD metrics were calculated from the fitted curve to further investigate how ITD tuning shapes depend on the auditory experience: best ITDs, derived from peak shaped responses, and ITD_MS,_ ITD at the maximum slope. Figure 5B compares the distribution of best ITDs across the three animal groups. ITD_best_ in the ED-CS group showed a statistically significant contralateral bias (two-tailed *t*-test, mean=276.6 µs, *t*(55)=5.6837, p<0.001), similarly as in AD group (two-tailed *t*-test, mean=162.1 µs, *t*(52)=3.4852, p=0.0010). In contrast, the distribution of ITD_best_ in ED-US group did not show a contralateral bias (two-tailed *t*-test, mean=91.7 µs, *t*(22)=0.9316, *p*=0.3616,). However, the difference in mean ITD_best_ between the three deafness groups did not reach statistical significance (*F*(2, 131)=2.39, *p*=0.0953, ANOVA). ITD_MS_ was broadly distributed with a mean near zero for all three groups (ED-CS: −53.4 µs, AD: −78.3 µs, ED-US: 8.2 µs) and there was no statistically significant difference between the three deafness groups (*F*(2, 262)=0.32, *p*=0.7261, ANOVA) (Figure 5C). In summary, there was minimal effect of different auditory experience during development on the distribution of ITD tuning shapes or the ITD tuning metrics except for the presence of a contralateral bias in the ITD_best_ in ED-CS group.

### Chronic CI stimulation did not improve temporal coding in early deaf animals

To characterize how the temporal response pattern to electrical stimulation depends on auditory experience, we measured responses to periodic electric pulse trains as a function of pulse rate. In all animal groups, there was a substantial variability in the pulse-rate dependence of firing rates and the relationship between responses during the on- and off-periods. Response patterns from the same three example neurons as in Figures 2 & 3 are shown in Figure 6. The neuron from the ED-US rabbit shows sustained and synchronized firing pattern up to 80 pps with higher firing rates than the firing rate during the off-period (excitatory response). For pulse rates above 80 pps, the response only lasts for ∼20 ms after the stimulus onset followed by a suppressive response for firing rates above 224 pps. The neuron from the ED-CS rabbit shows poorly synchronized response patterns with responses only lasting for 50-60 ms after stimulus onset followed by a suppressive period for pulse rates above 40 pps. Lastly the neuron from the AD rabbit showed sustained, synchronized response up to 112 pps. This neuron also shows small suppression of sustained firing rates for pulse rates above 112 pps.

**Figure 6.**
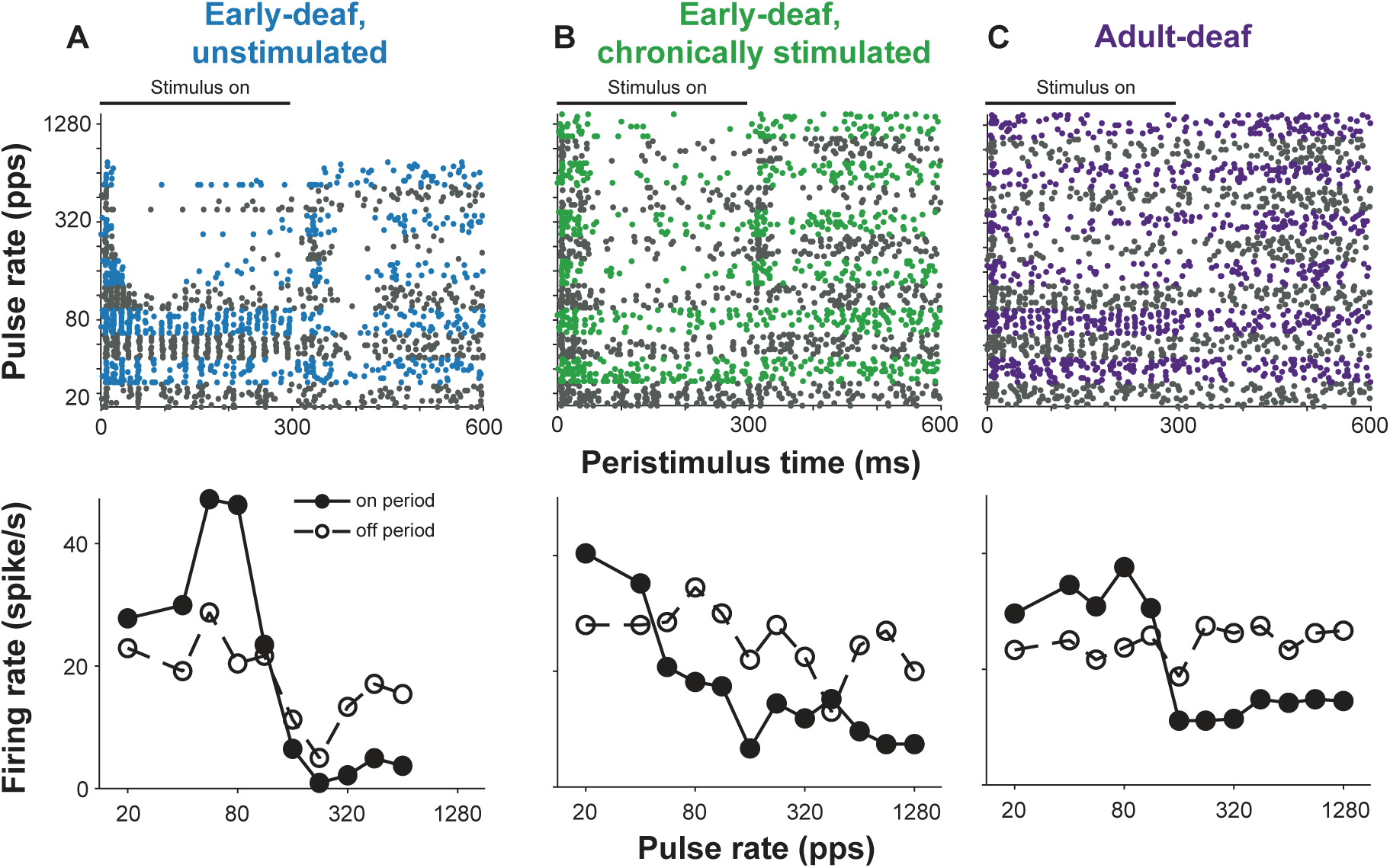
Top, Temporal discharge patterns (dot rasters) as a function of pulse rate for the three example neurons as in Figures 2 & 3. Alternating colors indicate blocks of stimulus trials at different pulse rate. Bottom, mean sustained firing rate as a function of pulse rate during stimulus on-period and-off period for the three example neurons.

For each pulse rate tested, the firing rate during the on-period (excluding the onset of 30 ms) and the off-period (excluding the offset of 100ms) were compared on a trial-by-trial basis to determine whether the periodic pulse train evoked significant excitatory or suppressive activity. Figure 7A shows the fraction of IC neurons showing excitatory and suppressive responses to pulse train stimuli as a function of pulse rate for the three deafness groups. Positive ordinates indicate the fraction of excitatory response and negative ordinates the fraction of suppressive responses. Overall there was a significant effect of pulse rate (*F*(11,35)=38.95, *p*<0.001) and deafness groups (*F*(2,35)=81.11, *p*<0.001) revealed by a two-way ANOVA on the arcsine-transformed fraction of excitatory units. Previously we found a general reduction in excitatory activity in the ED-US group compared to the AD group which was confirmed by post hoc paired comparisons (Tukey-Kramer). Surprisingly, we found a further reduction in excitatory activity in the ED-CS group compared to the ED-US group confirmed by post hoc paired comparisons (Tukey-Kramer, *p*<0.001). There was also an increase in suppressive activity in both early-deaf groups compared to the AD group. A two-way ANOVA on the arcsine-transformed fraction of suppressive units showed significant effects of both pulse rate (*F*(11,35)=12.08, *p*<0.001) and deafness groups (*F*(2,35)=35.49, *p*<0.001). A post hoc paired comparisons (Tukey-Kramer) confirmed that the fraction of suppressive neurons is significantly lower in the AD group compared to both ED groups (*p*<0.001 for both) but there was no difference between the two ED groups (*p*=0.8717).

**Figure 7.**
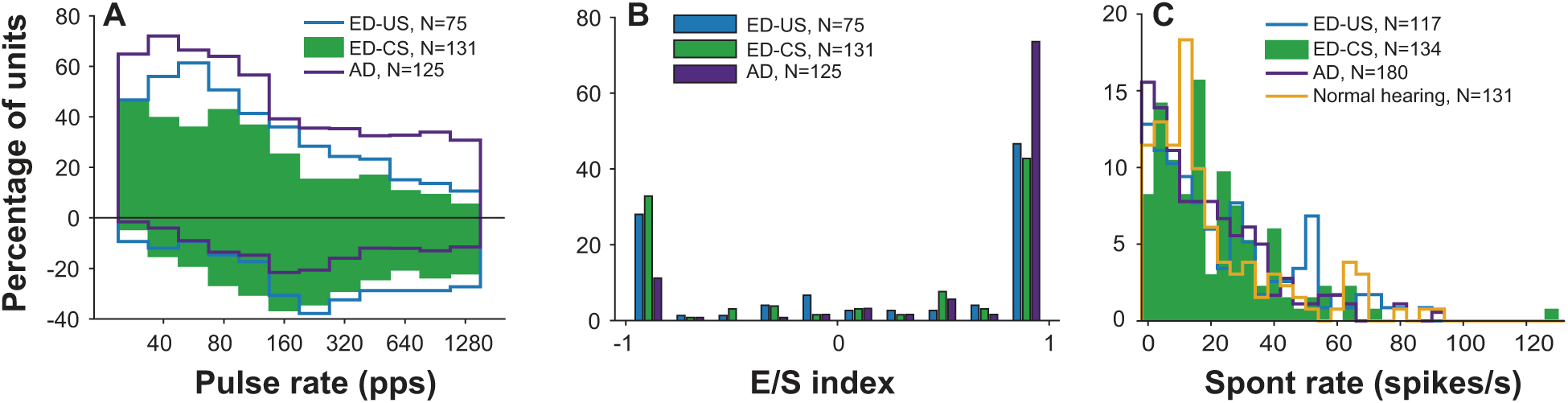
A, Percentage of units showing significant excitatory (positive ordinates) and suppressive (negative ordinates) response to pulse train stimulation as a function of pulse rate for the three animals groups. B, Distribution of E/S index for the animal groups. C, Distribution of spontaneous firing rates between the three animals groups and normal hearing animals.

We defined the excitatory/suppressive (E/S) ratio for each unit to characterize the relative prevalence of excitatory vs. suppressive responses across the pulse rates tested in the three animals groups (Chung et al. 2019) (Figure 7B). The E/S index ranges from −1 for purely suppressive response to +1 for purely excitatory response at all rates. A Kruskal-Wallis test showed a significant difference between deafness groups (χ^2^(2, 330)=18.82, *p*<0.001). Post hoc paired comparisons (Tukey-Kramer) showed that the median E/S ratio was larger in the AD group compared to both ED groups but there was no difference between the two ED groups. Similarly to the findings based on the fractions of excitatory and suppressive responses, chronic stimulation did not increase the reduced E/S ratio in early-onset deafness.

The difference in relative fractions of excitatory/suppressive response might be driven by differences in spontaneous activity of the IC neurons. We have previously shown that there was no significant difference in spontaneous firing rates in AD, ED-US and normal hearing animals (Chung et al., 2019). Figure 7C compares the distribution of SR in three deafness groups and normal hearing rabbits. Consistent with the previous results, the median spontaneous firing rates in the ED-CS group were in a similar range as the other deafness groups (ED-US=16.4 spikes/s, ED-CS=16.6 spikes/s, AD=15.1 spikes/s, NH=12.6 spikes/s) and did not differ statistically (Kruskal-Wallis test, χ^2^(3, 561)=3.73, *p*=0.2919). In general, we did not find an effect of deafness and different auditory experience on spontaneous activity.

We examined if there was an effect of chronic bilateral CI stimulation on synchronized activity to pulse train stimuli. For each neuron and pulse rate, the spike train was cross correlated to the stimulus to quantify the neural synchrony and define the upper pulse rate limit of synchronized activity (Hancock et al. 2012, Chung et al. 2014, 2019). Figure 8A compares the fraction of neurons that showed significant synchronized response as a function of pulse rate in the three deafness groups. For all pulse rates, the AD group shows the largest fraction of synchronized responses compared to two ED groups. Similar with the fraction of excitatory responses, the ED-CS group showed a smaller fraction of synchronized responses compared to the ED-US group for all rates tested. A two-way ANOVA on arcsine-transformed fractions shows significant effects of both deafness group (*F*(2, 35)=29.81, *p*<0.001) and pulse rate (*F*(11, 35)=107.12, *p*<0.001). Post hoc paired comparisons (Tukey-Kramer) show that the fraction of synchronized responses in the ED-US group is smaller than in the AD group (*p*=0.0033) but larger than in the ED-CS group (*p*=0.0024). Figure 8B compares the cumulative distribution of upper limit of synchronized response in three deafness groups. In this analysis, neurons that did not synchronize to any pulse rate were excluded. The cumulative distributions from three deafness groups nearly overlap with each other and did not differ statistically (Kruskal-Wallis test, χ^2^(2,256)=3.25, *p*=0.1973).

**Figure 8.**
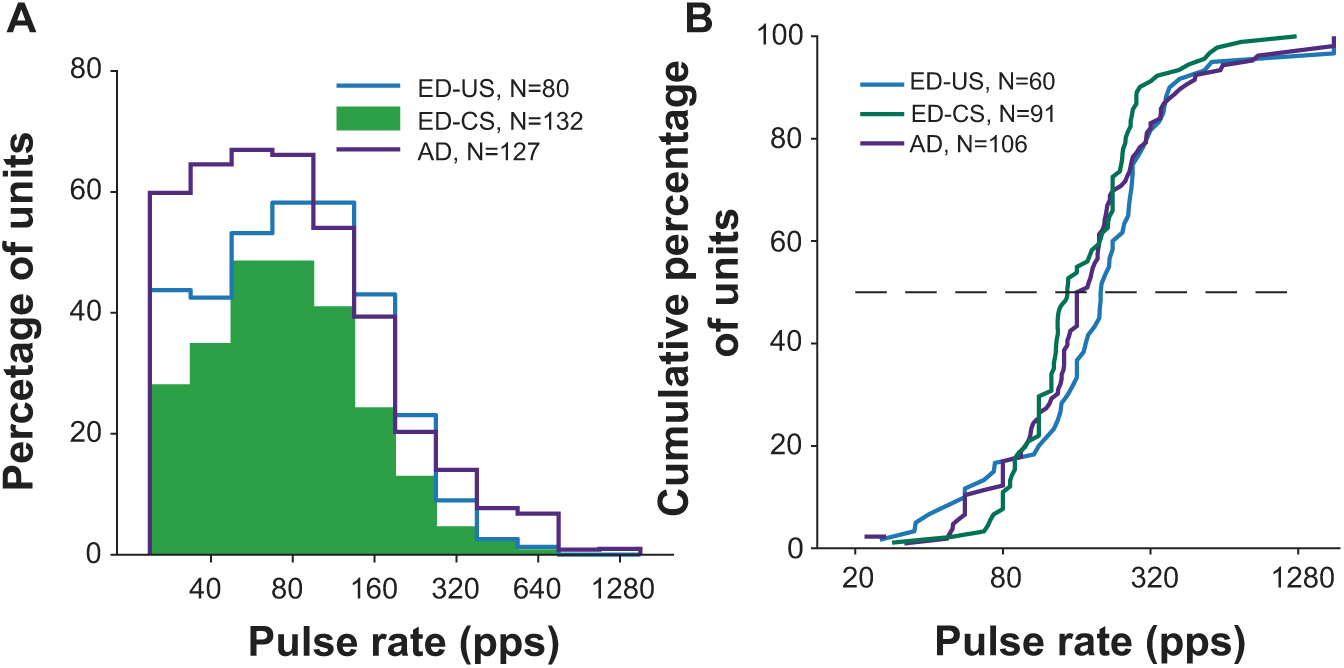
A, Percentage of units showing significant synchronized response to pulse train stimulation as a function of pulse rate for the three animal groups. B, Cumulative distribution of upper limits of synchronized response for the three animal groups.

In summary, chronic CI stimulation did not improve temporal coding in early deaf animals. Fewer neurons in the ED-CS group showed sustained excitatory response and synchronized response to pulse train stimulation. Therefore, the restoration of response properties with bilateral CI stimulation was limited to ITD sensitivity.

## Discussion

We characterized neural ITD sensitivity and temporal coding of IC neurons in rabbits that were neonatally deafened and chronically stimulated with bilateral CIs during development (ED-CS). The response properties were compared to data from both rabbits that were deafened as neonates and received no stimulation during development (ED-US) and rabbits with normal auditory development that were deafened as adults (AD). We found that chronic bilateral CI stimulation during development can reverse the effect of auditory deprivation and partly restore ITD sensitivity in the IC neurons. The prevalence of ITD sensitive neurons in the ED-CS rabbits was restored to a level comparable to AD rabbits. We also found improvement in ITD sensitivity measured by STVR in the ED-CS rabbits compared to ED-US rabbits; however, overall ITD sensitivity in the ED-CS rabbits was still poorer than in the AD rabbits. The improvement in prevalence and degree of neural ITD sensitivity was largest in the response to high stimulation rates. In contrast, we did not find an improvement in overall prevalence of excitatory response and synchronized response in ED-CS rabbits compared to ED-US rabbits. Rather, we found a decrease in the prevalence of excitatory responses and synchronized responses in the ED-CS group compared to the ED-US group. We did not find a difference in spontaneous activity in the ED-CS group compared to the ED-US and AD groups consistent with the previous results that showed the lack of an effect of auditory deprivation on spontaneous firing rate in the IC neuron in unanesthetized rabbits.

### Comparison with previous studies

Previous studies in anesthetized cats have reported improved temporal coding of electrical pulse train stimulation in the IC of animals that were neonatally deafened and chronically stimulated during development (Snyder, Leake, Rebscher, & Beitel, 1995; Vollmer et al., 1999) and also in neonatally-deafened animals that were given chronic stimulation after a long period of auditory deprivation (Vollmer, Leake, Beitel, Rebscher, & Snyder, 2005). In contrast, the current study did not find a reversal of effect of auditory deprivation in neural temporal coding.

This difference may stem from the difference in temporal properties of the chronic CI stimulation. We used the “Fundamental Asynchronous Stimulus Timing” (FAST) strategy that is designed to deliver ITD cues and employs lower average pulse rate than previous studies that found a restoration of temporal coding. Indeed, Vollmer et al. (1999) showed that the effect of passive stimulation on temporal coding in the IC neurons depends on the temporal properties of the electrical stimulation. Neonatally-deafened animals that received chronic low-rate stimulation (30-80 pps) did not show an increase in upper limit of synchronized response compared to unstimulated animals, while the group received high-rate stimulation (300 pps) showed an improvement in temporal coding. The lack of improvement in neural temporal coding associated with the chronic stimulation is likely due to the low average stimulation rate of FAST strategy.

The observation of an improvement in neural ITD sensitivity without an improvement in neural temporal coding in the auditory midbrain, raises an interesting question about the neural mechanism behind the limited temporal processing and ITD sensitivity in CI listening. The perceptual limit of ITD sensitivity at high pulse rates is similar in some respects to the limit of monaural rate discrimination in CI users. Most CI listeners can discriminate the rate of periodic pulse trains up to ∼300 pps (Moore & Carlyon, 2010). Similarly, most CI users show a degradation in perceptual ID sensitivity for pulse rates above ∼300 pps (Kan & Litovsky, 2015; Laback et al., 2015). Ihlefeld et al. (2015) showed a small correlation between monaural rate discrimination and ITD discrimination performance across different pairs of electrodes in the same subjects, suggesting that temporal coding and ITD sensitivity may dependent on a common neural mechanism. The current study shows that better neural ITD coding can be achieved without better temporal coding, suggesting there might be independent mechanisms behind the two processes. While temporal coding and ITD sensitivity are closely linked at the primary sites of binaural interaction in the medial and lateral superior olives (Batra & Yin, 2004), this is not the case in the IC, where ITD is coded in the average firing rate, while temporal coding may be limited by synaptic time constants in the IC itself.

### Mechanisms for effect of CI stimulation during development

The restoration of ITD sensitivity observed in IC neurons with chronic CI stimulation in early-deaf animals could be mediated by neural plasticity at any site along the ascending auditory pathway, just as the degradation in unstimulated animals could be caused by damage at any site. Although the reduction in spiral ganglion cell counts and demyelination of auditory nerve fibers can contribute to the degraded temporal coding in deafness (Hardie & Shepherd, 1999; Resnick, O’Brien, & Rubinstein, 2018), this effect is not limited to early-onset deafness (Shepherd & Javel, 1997) and is unlikely to be the mechanism behind degraded temporal and ITD coding in the midbrain being more prominent in early-onset deafness. Nevertheless, chronic CI stimulation can help prevent the degeneration of spiral ganglion cells in cats deafened by ototoxic drugs (Leake, Hradek, Rebscher, & Snyder, 1991; Leake, Hradek, & Snyder, 1999). Structural abnormalities in the endbulb synapses between auditory nerve fibers and spherical bushy cells of the cochlea nucleus in congenitally-deaf cats can be reversed by chronic CI stimulation during development (O’Neil, Limb, Baker, & Ryugo, 2010; Ryugo, Kretzmer, & Niparko, 2005). Because these synapses are specialized for transmission of precise temporal information, abnormalities in their morphology might degrade temporal and ITD processing in the medical superior olive (MSO). Tirko and Ryugo (2012) showed that the reduction in the size of excitatory synaptic boutons and in the number of inhibitory synapses onto MSO principal cells which occurs in the congenitally-deaf cats can be partly revered by chronic bilateral CI stimulation during development. The restorative effect of chronic CI stimulation in the current study was limited to ITD sensitivity, therefore it likely originates at sites that are specific for binaural processing.

### Implication for human performance and rehabilitation

Bilateral CIs have become the standard of care for children with congenital deafness. Children who received simultaneous bilateral CIs at an early age show partial reversal in the degraded cortical representation of ITDs due to deafness (Easwar et al., 2017). Similarly, bilateral CI stimulation starting at an early age can help improve perception of binaural cues, although not nearly to the level of normal hearing peers (Gordon et al., 2014). Results from the current study gives further evidence for the importance of appropriate auditory experience during development in early deafness. Our “chronic” stimulation only lasted 25 hours per week (5 hours/day, 5 days/week) yet achieved a significant therapeutic effect in restoring binaural sensitivity. Longer duration of stimulation with more meaningful behavioral context might result in even better outcomes.

Children in the studies cited above were provided with clinical bilateral CIs that are not designed to deliver ITD cues as the processors used in the current experiment. Currently clinical devices only provides envelope ITD cues. Whether further improvement in binaural sensitivity can be achieved with better access to ITD cues is unknown. Results from the current study alone do not show that precise ITD cues in the bilateral CI stimulation are necessary for the reversing the degradation in ITD sensitivity in early-onset deafness. However, Fallon et al. (2015) found no significant restorative effect of bilateral CI stimulation on binaural sensitivity with conventional continuous interleaved sampling (CIS) stimulation in IC neurons in anesthetized cats. On the other hand, our results also suggest that temporal properties of the electrical stimulation might be important for the development of temporal coding mechanism in the central auditory pathway. A processing strategy that can deliver precise binaural cues without a reduction in stimulation rate might be necessary for the restoration of both temporal and binaural sensitivity with CIs. Recent studies have shown that introduction of short inter-pulse intervals in high-rate pulse trains can enhance ITD sensitivity both in human perception (Srinivasan, Laback, Majdak, & Delgutte, 2018) and in IC neurons (Buechel, Hancock, Chung, & Delgutte, 2018).

In summary, we showed partial, but significant, restoration of neural ITD sensitivity in early-deaf animals with chronic bilateral stimulation. The restoration of ITD sensitivity was independent of neural temporal coding which did not show any improvement compared to unstimulated animals.

## Materials and Methods

### Animals and experiment design

All experiments were approved by the animal care and use committee of Massachusetts Eye and Ear. Four rabbits (three females, and one male) were neonatally deafened as neonates and implanted at 2 months of age. Chronic stimulation with wearable bilateral processors commenced 1–2 weeks after implantation. Neurophysiological data from the inferior colliculus was collected from all four implanted rabbits starting at 6 months of age and lasting for 3–7 months.

Neural data from these four early-deafened and chronically stimulated (ED-CS) rabbits are compared with reanalyzed data from four adult-deafened (AD) rabbits (Chung et al., 2016) and three early-deafened and unstimulated (ED-US) rabbits (Chung et al., 2019) used in previous studies. The AD rabbits were bilaterally deafened and implanted at 8-10 months of age and studied for ∼6 months. These rabbits were deafened during the implantation surgery by injection of distilled water into the scala tympani that causes hair cell death through osmotic stress (Chung et al., 2019; Chung et al., 2014). The three ED-US rabbits were deafened by daily injections of neomycin and implanted at 2-3 months of age as in the current study. Here, we only describe the methods in detail for the ED-CS group because the methods for other groups were reported in previous reports.

### Neonatal deafening

Newborn rabbits were deafened by daily subcutaneous injections of neomycin sulfate (60 mg/kg/day) from the first postnatal day (Chung et al., 2019). Auditory brainstem response (ABR) to acoustic clicks were measured at P22 under sedation (midazolam 2 mg/kg I.M., fentanyl 10 mcg/kg I.M.). In all neonatally-deafened rabbits, no measurable ABRs were confirmed at 110 dB SPL and the injections were discontinued.

### Surgical procedures

All surgical procedures were performed under anesthesia (xylazine 6 mg/kg S.C., ketamine 35 mg/kg I.M. then maintained by isoflurane 2.5% delivered via facemask in O_2_ 0.8 l/min) and strict aseptic conditions.

At 2 months of age, the rabbits were implanted with intracochlear electrode arrays bilaterally. The method of cochlear implantation was the same as described in detail previously (Chung et al. 2014, 2016, and, 2019). In brief, the bulla was exposed via the retroauricular approach, and opened with a bone drill to visualize the basal turn of the cochlea and round window. The round window was drilled and enlarged enough to achieve full insertion of an HL-8 eight contact electrode array (Cochlear Ltd.) without resistance. The opening of the enlarged round window and bulla were sealed carefully with a small piece of muscle and bone cement, respectively. CI connectors were attached on the skull with stainless steel screws and dental cement. Immediately after the CI surgery, postoperative imaging, including skull X-ray and computed tomography, was performed to detect possible electrode misplacements and to assess insertion depth. Electrode misplacement such as tip-over, loop, kinking, and scalar crossing was not observed in any cases.

At 6 months of age, a brass bar was attached to the skull and a cylindrical barrier (1.5–2 cm diameter) for a sterile portal to a subsequent craniotomy was built with dental cement. After recovery from headpost surgery (∼1 week), rabbits were habituated to the experimental setup until they could sit quietly for 2–3 hours with the head fixed while receiving electrical stimulation. After the habituation period (1–2 weeks), a third surgery was performed to permit insertion of a recording electrode into the inferior colliculus (IC). Two small craniotomies were made, one on each side of the parietal bone inside the cylindrical barrier. The exposed dura of the occipital cortex was covered with topical antibiotic ointment and the cylinder was sealed with dental impression material. Neurophysiological recordings through the craniotomy commenced after 3 days of recovery.

### Chronic stimulation with bilateral wearable processors

One week after the implant surgery, the CIs were activated. The programming process was performed in awake state using the Custom-Sound programming software (Cochlear Ltd.). At the first step in programming, impedance and electrically evoked compound action potential (ECAP) for each intracochlear electrode contact were measured to assess the effectiveness of cochlear implantation. Electrodes with abnormally high impedance and shorted electrodes were deactivated and not used for further study. Electrode impedances and ECAP thresholds, as indicators of physiologic and implant function, were measured weekly and monthly, respectively. For the next step of programming, the number of activated channels and electrode pairs between ears were matched based on electrical measures and radiographic results. In two animals with the same depth of electrode insertion in both ears, the most basal electrode was not activated due to high ECAP threshold, and the apical seven electrodes were used. In two other animals with asymmetric insertion depth, only six pairs of matched electrodes were stimulated. The ECAP threshold profile and animal’s behavioral response was used to estimate the upper-stimulation levels and expected threshold levels assuming a typical electrical dynamic range (6.3 dB as 40 current levels in Nucleus devices).

We used an experimental “Fundamental Asynchronous Stimulus Timing” (FAST) strategy specifically designed to deliver ITD cues (Smith, 2010, 2014) to stimulate the cochleas. Chronic environmental stimulation commenced approximately 2 weeks after CI surgery. Passive stimulation with environmental sounds associated with normal activities and sounds of nature played via loudspeaker took place in the rabbits’ home cage in the animal care facility for up to 5 hours per day, 5 days per week. During the stimulation sessions, rabbits wore the modified CI device pair (Nucleus CP800 sound processor and CI emulator T51107, Cochlear Ltd.) in a backpack with the outside auxiliary microphones positioned posterior to the ipsilateral ear.

In addition to the passive environmental stimulation, active behavioral training by operant conditioning was performed 3 days per week, for up to 3 hours per day, in a sound-treated booth where the rabbit was able to move freely. The task was to find the sound target using auditory cues in a rectangular area. The rabbits wore the CI processors connected with Bluetooth audio receiver within the sound processor backpack. During behavioral sessions, the animal was monitored via a video camera and the distance and head orientation in relation to the target was achieved by a computer to present stimuli with the relevant binaural cues under direct computer control. 100 Hz pulse train were delivered to a pair of monopolar channels with interaural time difference (ITD), interaural level difference (ILD), and mean level calculated based on rabbit’s relative location and head orientation from the target.

Initially, the rabbits were trained to find a sound target using auditory cues, as well as visual stimuli from light emitting diodes. When the rabbits learned and performed the task with an accuracy above chance level, the rabbits were weaned on visual stimuli and trained with auditory stimuli alone. Maximum four target food bowls with pellet feeders were placed at each corner of a rectangular arena. The stimuli on each trial were initiated when the location of sound target was randomly selected (except the nearest one to the animal) by a computer. A food pellet was given as a reward if the rabbit successfully reached the correct target within 60 seconds. Reaching the incorrect target or standing out of any targets over one minute, which was determined as failed trial, discontinued the stimuli. Light in the booth turned off for 10 seconds in failed trials. Because none of the rabbits showed any significant progress in performance with auditory stimuli alone after 40 sessions, the behavioral training sessions were discontinued after 3 months.

### Electrophysiological methods

#### Single-unit recordings

All neurophysiological recording procedures were the same in three deafness groups. Polyimide insulated platinum/iridium linear microelectrode arrays with 4–6 contacts (150 µm spacing between contacts, 12.5 µm site diameter; MicroProbes) were used to record single-unit activity from the IC. Recording electrodes were advanced in a dorsoventral direction through the occipital cortex down to the IC, which was identified by entrained multi-unit activity to the search stimuli. We sampled neurons across all depths where this background entrainment was observed. The point of entry for the electrode was varied within the craniotomy to widely sample the IC.

To reduce the electrical stimulus artifact, recordings were made differentially between one contact showing clear spike activity and a local reference obtained by averaging the signals from the remaining contacts. Signals from the recording electrodes were acquired by a unity gain headstage (HST/16o50; Plexon) and then filtered (100–8000 Hz) and amplified (PBX2; Plexon). The conditioned signals were sampled at 100 kHz using a 12-bit A/D converter. Then the stimulus artifact was removed by a gate-and-interpolate technique (Heffer & Fallon, 2008) for low pulse rates (< 200 pps). For measurements of ITD sensitivity at higher pulse rates, an offline artifact rejection method based on template subtraction was used (Buechel et al., 2018).

#### Stimuli

All stimuli were trains of biphasic pulses (50 µs/phase) delivered to each cochlear implant through a pair of custom-built, high-bandwidth, isolated current source. Current was passed between used the most apical and basal electrode sites which excites neurons over a wide range of tonotopic axis. The search stimulus was a sequence of three pulses presented successively to both ear (diotically), the left ear and the right ear with a 100-ms interval between each pulse 200-ms silent interval between the triplets. Upon isolating a single unit, the firing rate was obtained as a function of current level to determine the response threshold and the dynamic range. Spontaneous firing rate was measured by recording firing rate during 30-s of silence. All units showing a driven response to the search stimuli, whether excitatory or suppressive, were further studied and included in the database.

To characterize neural temporal coding, responses to diotic pulse trains with pulse rates ranging from 20–1280 pps in half-octave steps with 12 repetitions were obtained. Stimuli were usually presented at 1–6 dB above the threshold of synchronized responses to a single diotic pulse.

ITD sensitivity was characterized by acquiring responses while varying the relative timing of pulse trains delivered to the two implants with zero interaural level difference (ILD). We first obtained response by varying ITDs from −1500 to +1500 µs in 300-µs steps at 80 pps for three to four current levels within the dynamic range of the unit. The current level that yielding the strongest ITD sensitivity was selected to further investigate the effect of varying pulse rate. ITD was then varied from −2000 to +2000 μs in 200 μs steps with 10 repetitions at each ITD. To investigate the effect of pulse rate on ITD sensitivity, pulse rates ranging from 20 to 640 pps in one octave steps were randomly interleaved with the ITD values tested, resulting in a matrix of firing rates. For both measurements of temporal coding and ITD tuning, the stimulus duration was 300 ms with silent interval of 300 ms, resulting in repetition period of 600 ms.

### Statistical Analysis

#### Neural ITD sensitivity

ITD tuning curves were obtained by averaging the firing rates for each ITD over the entire stimulus duration (0–300 ms) and across all stimulus presentations. Neural sensitivity to ITD was quantified by the ITD signal-to-variance ratio (STVR) (Chung et al., 2016; Hancock et al., 2010). ITD STVR is an ANOVA-based metric that represents the fraction of variance in firing rates due to changes in ITD relative to the total variance in firing rate across both stimulus trials and ITDs. Responses were considered to be ITD sensitive when the STVR was significantly greater than zero (F test, p < 0.01). The degrees of freedom for the F test were (20,189) for most cases when 21 ITD values were tested over 10 repetitions.

The ITD tuning curves were fitted to a template consisting of the sum of Gaussian and sigmoid function to classify the shape (Chung et al., 2019; Chung et al., 2016):

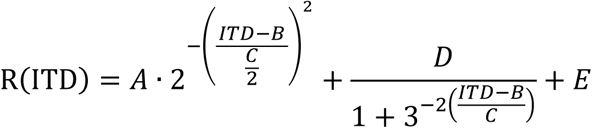

where A, D, and E are scaling factors, B is the center of both Gaussian and the sigmoid functions, and C represents the half-width of the Gaussian and the half-rise of the sigmoid function. The fitted curves were classified into four types: monotonic, peak, trough, and unclassified (ITD curves that were poorly fit by the template, *r*^2^<0.75). From the fitted curves, the best ITD (ITD_best_), ITD at the maximum of peak shaped responses, and ITD_MS,_ ITD at the maximum slope, were derived. Only ITD tuning curves in response to pulse rates below 160 pps were used in shape classification and to calculate of ITD metrics to avoid multiple cycles of periodicity within the measured ITD range of ±2000 µs.

#### Excitatory and suppressive responses

To quantify the prevalence of excitatory and suppressive response in different deafness groups, the “sustained” firing rate during the pulse train presentation (“on-period”) was compared to the sustained firing rate during the silent interval between the stimuli (“off-period”). The first 30 ms of the 300 ms stimulus was excluded to isolate sustained, on-period response from the onset response. Likewise the first 100 ms of the 300 ms interstimulus period to isolate sustained, off-period response from rebound response.

The prevalence of sustained excitatory and suppressive responses was quantified by two methods: 1) comparing the firing rates during the on-period and off-period on a trial-by-trial basis and 2) excitatory/suppressive (E/S) index to quantify the relative difference in excitatory and suppressive activity across pulse rates for each unit (Chung et al., 2019). If the on-period mean firing rate was significantly higher (resp. lower) than the off-period mean firing rate (paired t-test, two-tailed, p<0.01), the response was considered excitatory (resp. suppressive). E/S index is obtained by comparing the excitatory area E and suppressive area A. The excitatory area E was defined as the sum of the on-period firing rates minus the off-period firing rates over pulse rates that evoked a significant excitatory response. Similarly, the suppressive area S was defined as the sum of the off-period firing rate minus the on-period firing rate over pulse rates that evoked a significant suppressive response. The E/S index was the ratio (E-S)/(E+S) that ranges from −1 for purely suppressive response to +1 for purely excitatory response.

#### Synchronized response to pulse trains

The prevalence of synchronized response and the upper limit of pulse rate that can elicit synchronized response were compared across deafness groups. To quantify the degree of synchrony of spike to the pulse trains, the stimulus pulse train was cross-correlated with the spike trains (Chung et al., 2019; Chung et al., 2014; Hancock, Chung, & Delgutte, 2012). The 30-ms onset response was excluded from the spike train to avoid biasing the synchrony estimate by prominent peaks in the onset response. To assess the statistical significance of cross-correlation peaks, the cross-correlation was computed for 5000 random spike trains containing the same number of spikes as in the original neural recording. A confidence bound for the null hypothesis of no correlation was defined as the 99.5th percentile of the cross-correlograms across the 5000 random spike trains. A peak in the cross-correlogram was considered significant when it exceeded the confidence bound. The upper rate limit of synchronized responses was computed by linearly interpolating the height of the cross-correlogram peaks to find the pulse rate where the peak height intercepted the confidence bound.

### Histological processing

In three early-deaf and chronically stimulated rabbits, we made electrolytic lesions during the last recording session while the animal was under anesthesia (xylazine, 6 mg/kg, S.C.; ketamine, 44 mg/kg, I.M.). Electrolytic lesions were made by passing radiofrequency current for 60 seconds supplied through a lesion maker (Model LM-3, Grass Instruments) to mark the borders of the region showing evoked activity with CI stimulation. The rabbit was then perfused intracardially using a 10% formalin solution. The brain was immersed in fixative for 24 h and then transferred to 25% sucrose solution for several days. The brain was embedded in optimal cutting temperature (OCT) compound and sagittal sections (80 μm) were cut with a cryostat at −15°C. Sections were mounted on gel-subbed slides and dehydrated with ethanol bath of increasing concentration up to 100%. Cell bodies were stained with azure-thionin. Electrode traces reaching the central nucleus of the IC were identified as they passed through the occipital cortex and superior colliculus. All lesions were located in the central nucleus of the IC.

## Acknowledgements

This work was supported by the National Institutes of Health Grant R01 DC005775. We thank Ken Hancock, Ishmael Stefanov-Wagner, Alice Gelman, Joseph Wagner, Stephanie Ventura and Marie Ortega for technical assistance. We thank Hugh Curtin, Renee Mitchell for assistance in radiographic imaging and Kevin Franck for the advice on programming of the cochlear implant processors. Lastly, we thank Zachary Smith at Cochlear Ltd for providing the cochlear implant sound processors and software.

## References

Batra, R., & Yin, T. C. (2004). Cross correlation by neurons of the medial superior olive: a reexamination. J Assoc Res Otolaryngol, 5(3), 238–252. doi:10.1007/s10162-004-4027-4

Buechel, B. D., Hancock, K. E., Chung, Y., & Delgutte, B. (2018). Improved Neural Coding of ITD with Bilateral Cochlear Implants by Introducing Short Inter-pulse Intervals. J Assoc Res Otolaryngol, 19(6), 681–702. doi:10.1007/s10162-018-00693-0

Chung, Y., Buechel, B. D., Sunwoo, W., Wagner, J. D., & Delgutte, B. (2019). Neural ITD Sensitivity and Temporal Coding with Cochlear Implants in an Animal Model of Early-Onset Deafness. J Assoc Res Otolaryngol, 20(1), 37–56. doi:10.1007/s10162-018-00708-w

Chung, Y., Hancock, K. E., & Delgutte, B. (2016). Neural Coding of Interaural Time Differences with Bilateral Cochlear Implants in Unanesthetized Rabbits. The Journal of neuroscience, 36(20), 5520–5531. doi:10.1523/JNEUROSCI.3795-15.2016

Chung, Y., Hancock, K. E., Nam, S. I., & Delgutte, B. (2014). Coding of electric pulse trains presented through cochlear implants in the auditory midbrain of awake rabbit: comparison with anesthetized preparations. The Journal of neuroscience, 34(1), 218–231. doi:10.1523/JNEUROSCI.2084-13.2014

Easwar, V., Yamazaki, H., Deighton, M., Papsin, B., & Gordon, K. (2017). Cortical Representation of Interaural Time Difference Is Impaired by Deafness in Development: Evidence from Children with Early Long-term Access to Sound through Bilateral Cochlear Implants Provided Simultaneously. J Neurosci, 37(9), 2349–2361. doi:10.1523/JNEUROSCI.2538-16.2017

Fallon, J., Irving, S., Wise, A., & Irvine, D. R. (2015). Effects of cochlear implant use on binaural processing. Paper presented at the Assoc Res Otolaryngol Abstr.

Gordon, K. A., Deighton, M. R., Abbasalipour, P., & Papsin, B. C. (2014). Perception of binaural cues develops in children who are deaf through bilateral cochlear implantation. PLoS One, 9(12), e114841. doi:10.1371/journal.pone.0114841

Hancock, K. E., Chung, Y., & Delgutte, B. (2012). Neural ITD coding with bilateral cochlear implants: effect of binaurally coherent jitter. Journal of neurophysiology, 108(3), 714–728. doi:10.1152/jn.00269.2012

Hancock, K. E., Noel, V., Ryugo, D. K., & Delgutte, B. (2010). Neural coding of interaural time differences with bilateral cochlear implants: effects of congenital deafness. The Journal of neuroscience, 30(42), 14068–14079. doi:30/42/14068 [pii] 10.1523/JNEUROSCI.3213-10.2010

Hardie, N. A., & Shepherd, R. K. (1999). Sensorineural hearing loss during development: morphological and physiological response of the cochlea and auditory brainstem. Hear Res, 128(1-2), 147–165.

Heffer, L. F., & Fallon, J. B. (2008). A novel stimulus artifact removal technique for high-rate electrical stimulation. J Neurosci Methods, 170(2), 277–284. doi:S0165-0270(08)00079-4 [pii] 10.1016/j.jneumeth.2008.01.023

Ihlefeld, A., Carlyon, R. P., Kan, A., Churchill, T. H., & Litovsky, R. Y. (2015). Limitations on Monaural and Binaural Temporal Processing in Bilateral Cochlear Implant Listeners. Journal of the Association for Research in Otolaryngology: JARO, 16(5), 641–652. doi:10.1007/s10162-015-0527-7

Kan, A., & Litovsky, R. Y. (2015). Binaural hearing with electrical stimulation. Hearing Research, 322, 127–137. doi:10.1016/j.heares.2014.08.005

Laback, B., Egger, K., & Majdak, P. (2015). Perception and coding of interaural time differences with bilateral cochlear implants. Hearing Research, 322, 138–150. doi:10.1016/j.heares.2014.10.004

Leake, P. A., Hradek, G. T., Rebscher, S. J., & Snyder, R. L. (1991). Chronic intracochlear electrical stimulation induces selective survival of spiral ganglion neurons in neonatally deafened cats. Hearing Research, 54(2), 251–271.

Leake, P. A., Hradek, G. T., & Snyder, R. L. (1999). Chronic electrical stimulation by a cochlear implant promotes survival of spiral ganglion neurons after neonatal deafness. J Comp Neurol, 412(4), 543–562.

Litovsky, R. Y., Jones, G. L., Agrawal, S., & van Hoesel, R. (2010). Effect of age at onset of deafness on binaural sensitivity in electric hearing in humans. J Acoust Soc Am, 127(1), 400–414. doi:10.1121/1.3257546

Litovsky, R. Y., Parkinson, A., & Arcaroli, J. (2009). Spatial hearing and speech intelligibility in bilateral cochlear implant users. Ear Hear, 30(4), 419–431. doi:10.1097/AUD.0b013e3181a165be

Moore, B. C. J., & Carlyon, R. P. (2010). Perception of Pitch by People with Cochlear Hearing Loss and by Cochlear Implant Users. In C. J. Plack, A. J. Oxenham, & R. R. Fay (Eds.), Pitch: Neural Coding and Perception (pp. 234–277): Springer.

Murphy, J., & O’Donoghue, G. (2007). Bilateral cochlear implantation: an evidence-based medicine evaluation. Laryngoscope, 117(8), 1412–1418.

O’Neil, J. N., Limb, C. J., Baker, C. A., & Ryugo, D. K. (2010). Bilateral effects of unilateral cochlear implantation in congenitally deaf cats. J Comp Neurol, 518(12), 2382–2404. doi:10.1002/cne.22339

Resnick, J. M., O’Brien, G. E., & Rubinstein, J. T. (2018). Simulated auditory nerve axon demyelination alters sensitivity and response timing to extracellular stimulation. Hear Res, 361, 121–137. doi:10.1016/j.heares.2018.01.014

Ryugo, D. K., Kretzmer, E. A., & Niparko, J. K. (2005). Restoration of auditory nerve synapses in cats by cochlear implants. Science, 310(5753), 1490–1492. doi:10.1126/science.1119419

Salloum, C. A., Valero, J., Wong, D. D., Papsin, B. C., van Hoesel, R., & Gordon, K. A. (2010). Lateralization of interimplant timing and level differences in children who use bilateral cochlear implants. Ear Hear, 31(4), 441–456. doi:10.1097/AUD.0b013e3181d4f228

Shepherd, R. K., & Javel, E. (1997). Electrical stimulation of the auditory nerve. I. Correlation of physiological responses with cochlear status. Hear Res, 108(1-2), 112–144.

Smith, Z. M. (2010). Improved sensitivity to interaural time differences with the FAST coding strategy. Paper presented at the 11th International Conference on Cochlear Implants and Other Implantable Auditory Technologies Stockholm, Sweden.

Smith, Z. M. (2014). Hearing better with interaural time differences and bilateral cochlear implants. The Journal of the Acoustical Society of America, 135(4), 2190–2191.

Snyder, R., Leake, P., Rebscher, S., & Beitel, R. (1995). Temporal resolution of neurons in cat inferior colliculus to intracochlear electrical stimulation: effects of neonatal deafening and chronic stimulation. J. Neurophysiol., 73(2), 449–467.

Srinivasan, S., Laback, B., Majdak, P., & Delgutte, B. (2018). Introducing Short Interpulse Intervals in High-Rate Pulse Trains Enhances Binaural Timing Sensitivity in Electric Hearing. J Assoc Res Otolaryngol, 19(3), 301–315. doi:10.1007/s10162-018-0659-7

Tillein, J., Hubka, P., Syed, E., Hartmann, R., Engel, A. K., & Kral, A. (2010). Cortical representation of interaural time difference in congenital deafness. Cereb Cortex, 20(2), 492–506. doi:bhp222 [pii] 10.1093/cercor/bhp222

Tirko, N. N., & Ryugo, D. K. (2012). Synaptic plasticity in the medial superior olive of hearing, deaf, and cochlear-implanted cats. J Comp Neurol. doi:10.1002/cne.23038

van Hoesel, R. J., Jones, G. L., & Litovsky, R. Y. (2009). Interaural time-delay sensitivity in bilateral cochlear implant users: effects of pulse rate, modulation rate, and place of stimulation. Journal of the Association for Research in Otolaryngology: JARO, 10(4), 557–567. doi:10.1007/s10162-009-0175-x

van Hoesel, R. J., & Tyler, R. S. (2003). Speech perception, localization, and lateralization with bilateral cochlear implants. The Journal of the Acoustical Society of America, 113(3), 1617–1630.

Vollmer, M., Leake, P. A., Beitel, R. E., Rebscher, S. J., & Snyder, R. L. (2005). Degradation of temporal resolution in the auditory midbrain after prolonged deafness is reversed by electrical stimulation of the cochlea. J Neurophysiol, 93(6), 3339–3355. doi:00900.2004 [pii] 10.1152/jn.00900.2004

Vollmer, M., Snyder, R. L., Leake, P. A., Beitel, R. E., Moore, C. M., & Rebscher, S. J. (1999). Temporal properties of chronic cochlear electrical stimulation determine temporal resolution of neurons in cat inferior colliculus. J Neurophysiol, 82(6), 2883–2902.

